# Space Use and Drivers of Daily Movement in the Common Crane *Grus grus* Wintering in the Agricultural Landscape of Western India

**DOI:** 10.64898/2026.05.17.725127

**Authors:** Harindra L. Baraiya, Anju Baroth, R. Suresh Kumar

## Abstract

Wintering migratory birds must balance energetic requirements, resource availability, and disturbance in increasingly human-modified landscapes. However, individual-level variability in daily movement and winter space use remains poorly understood in populations of the Common Crane wintering in South Asia. We investigated how seasonal dynamics, landscape composition, and individual differences structure winter movement ecology in a semi-arid agro-wetland system in western India. We analysed high-resolution GPS telemetry data from four tagged cranes tracked across three consecutive winters. Daily movement distances were modelled using mixed-effects approaches to partition variance within and among individuals and among winters. Daily movement trajectories were evaluated using non-linear temporal terms. Landscape predictors, including the proportion of cropland, built-up area, and habitat heterogeneity, were incorporated to assess environmental drivers. Winter utilisation distributions were estimated using dynamic Brownian bridge movement models. Daily movement distance followed a non-linear seasonal pattern consistent with shifts in the profitability of agricultural resources over winter. Cropland proportion and landscape evenness were negatively associated with movement distance, whereas a high proportion of built-up areas increased daily movement distance, reflecting a trade-off between resource concentration and anthropogenic disturbance. Winter utilisation distribution size varied markedly both within individuals across years and among individuals within seasons. Winter movement and space use in Common Cranes are predominantly context-dependent and environmentally driven. Seasonal dynamics, agricultural landscapes, and human disturbance jointly structure movement patterns, with limited but consistent individual differences. While limited by sample size, this multi-year, individual-based telemetry provides insight into winter spatial strategies of Common Cranes in dynamic semi-arid agro-wetland systems.

## INTRODUCTION

Movement is a fundamental component of animal ecology, linking individual behaviour to energetic balance, fitness, and broader population and community processes (Nathan *et al*. 2008, Morales *et al*. 2010). At fine temporal and spatial resolutions, movement patterns emerge from behavioural decisions shaped by interactions between internal states and external environmental conditions. The decision as to when and where to move is mediated by physiological requirements, energetic balance, and behavioural motivations, such as foraging, resting, or avoiding adverse conditions, together with abiotic and biotic factors including resource availability, microclimate, and predation risk (Nathan *et al*. 2008). Because movement directly determines both energy acquisition and energetic expenditure, fine-scale movement decisions both reflect and regulate bioenergetic trade-offs (Soriano-Redondo *et al*. 2023, Schindler *et al*. 2024, Cordes *et al*. 2025). Understanding the mechanisms underlying daily movement behaviour is therefore essential for understanding how individuals respond to spatial and temporal variation in their environments.

Human activities have transformed a significant portion of the Earth’s surface, altering the structure, connectivity, and disturbance regimes of terrestrial landscapes (Barnosky *et al*. 2012). The expansion of agriculture, urbanisation, and associated infrastructure has altered habitat configuration and introduced novel threats (Haddad *et al*. 2015, White & Razgour 2020, Cao & Song 2022, Roy *et al*. 2022). These changes influence landscape permeability, spatial distribution of resources and risk gradients, thereby shaping the movement decisions of animals inhabiting human-dominated landscapes (Avgar *et al*. 2016, Doherty *et al*. 2021, Therkildsen *et al*. 2021, Valle *et al*. 2024, Teitelbaum *et al*. 2026). Agricultural intensification is a major driver of land-use and land-cover (LULC) change and, together with habitat loss, is widely recognised as a leading cause of global wildlife declines, particularly among bird populations (Beckmann *et al*. 2019, Rigal *et al*. 2023, Das *et al*. 2025).

In agrarian landscapes, cropping systems introduce pronounced spatiotemporal variability through planting, growth, harvesting, and fallow cycles, creating shifting patterns of food availability within human-dominated environments (Butler *et al*. 2010, Vasseur *et al*. 2013). While agricultural intensification is widely associated with declines in avian populations, productive crops within agricultural landscapes can also provide highly attractive foraging resources for many bird species, including cranes and waterfowl, particularly geese (Fox *et al*. 2005, Nilsson *et al*. 2019, Buitendijk *et al*. 2022). Therefore, these species make a particularly informative model for studying the interaction between large grazing birds and agrarian landscapes through the lens of movement ecology [eg (Rosin *et al*. 2012, Nilsson *et al*. 2018, Wu *et al*. 2020, Overton & Casazza 2023, Clausen *et al*. 2024, Patterson *et al*. 2026, Teitelbaum *et al*. 2026)]. Cranes (family Gruidae) are long-lived, migratory species with high energetic demands and strong reliance on open habitats for foraging and roosting (Johnsgard 1983). Many crane populations have increasingly incorporated agricultural fields into their foraging strategies, exploiting post-harvest residues and cultivated crops as predictable and energy-rich food sources (Hemminger *et al*. 2022). As a consequence, cranes frequently occupy landscapes that are intensively managed and subject to rapid temporal change. Their daily movement patterns, therefore, reflect both the distribution of anthropogenic food resources and the spatial configuration of roosting and foraging habitats (Nilsson *et al*. 2018, 2020), making cranes well suited for investigating how large-bodied, avian herbivores respond to dynamic agrarian environments.

In western India, the common crane (*Grus grus*) arrives from northern Asia (Higuchi *et al*. 2008, Ram *et al*. 2023) to winter in heterogeneous agrarian landscapes that include croplands, rangelands, wetlands, and human settlements (Khachar *et al*. 1987). Across the Asian range, studies on wintering common cranes are geographically limited, largely concentrated in East Asia, and primarily focus on habitat use and behavioural ecology (Kong *et al*. 2018, Jia *et al*. 2019, Wang *et al*. 2019, Ming-Ming *et al*. 2021). On the Yunnan–Guizhou Plateau and in the Caohai Wetland, cranes exhibited strong diurnal routines, extensive daily movements between roosting and foraging sites, and behavioural segregation from sympatric crane species, likely facilitating coexistence through spatial and temporal niche partitioning (Kong *et al*. 2018, Ming-Ming *et al*. 2021). A broader wetland study from the Yangtze River floodplain further highlights the importance of water level, food availability, and distance from human settlements in shaping winter habitat selection and distribution patterns (Wang *et al*. 2019). These studies emphasise the ecological importance of agro-wetland mosaics, daily activity rhythms, and interspecific interactions, but offer limited insights into individual-level movement variability from high-resolution tracking.

GPS telemetry-based research has recently provided the first quantitative insights into winter range distributions, migration connectivity, and broad habitat associations of common cranes in India, highlighting a strong reliance on croplands and water bodies as well as a potential vulnerability to agricultural change and wetland loss (Ram *et al*. 2023). Earlier descriptive studies from Gujarat and Kutch documented the winter distribution, food habits, crop use, and roosting ecology, emphasising a strong dependence on agricultural landscapes and grassland resources, along with early warnings regarding habitat transformation and land-use change (Khachar *et al*. 1987, Tiwari & Rahmani 1996). Despite these foundational contributions, empirical understanding of the wintering ecology of the Common Crane in India remains largely descriptive and coarse in scale, with limited mechanistic insight into behavioural processes within wintering grounds. In particular, how fine-scale daily movement behaviour responds to landscape composition, habitat heterogeneity, and anthropogenic pressures in intensively cultivated agro-wetland systems remains poorly understood. Foraging behaviour has a major influence on individual fitness by shaping energy intake, locomotor costs, and exposure to risk. Examining movement patterns is therefore critical for understanding the energetic mechanisms that ultimately influence survival and reproductive success (Cordes *et al*. 2025). Because daily distance travelled reflects the spatial scale at which individuals balance resource acquisition against energetic expenditure and disturbance avoidance, it provides a mechanistic proxy for these trade-offs. However, quantitative analyses linking day-to-day movement distances with spatial variation in resource distribution and disturbance gradients are scarce in the region. These limitations constrain our ability to predict the behavioural responses of wintering common cranes to rapidly changing agricultural environments and to develop evidence-based conservation strategies for Asian populations that winter in India.

Here, we use high-resolution GPS tracking data from four common cranes over three wintering seasons in western India to investigate the landscape- and individual-level mechanisms that shape daily movement behaviour in an agrarian winterscape. We hypothesise that wintering movements are structured by winter progression, landscape composition, and individual behavioural strategies. Specifically, we predict that daily movement distances exhibit non-linear seasonal dynamics, decreasing in resource-rich landscapes and increasing in fragmented or disturbance-prone environments. We further expect that the spatial extent of winter utilisation distribution varies significantly among years. To test these predictions, we (ii) assessed the relative influence of landscape composition and winter progression on movement and (iii) estimated utilisation distribution separately for each winter to assess the interannual variation. Our study provides a primary, mechanistic, individual-based perspective on how common cranes adjust their daily movements in response to dynamic agrarian environments. More broadly, this work advances understanding of the movement ecology of large migratory birds in human-dominated winterscapes and highlights the role of individual-level variation in shaping movement patterns in modified landscapes.

## METHODS

### GPS tracking data and pre-processing

We obtained Movement data from GPS-GSM transmitters deployed on five adult common cranes and covered three wintering seasons (October to March) between 2022 and 2025 (Supplementary Table S1). We received permits to capture and tag common cranes from the office of the Principal Chief Conservator of Forest, Gujarat State Forest Department, vide letters WLP/ACF/RTC-3/C/1050-51/2018-19, dated 04-02-2019, and WL/RTC/28/C/852-853/2021-22, dated 19-01-2022. GPS locations were recorded at 10-60-minute intervals and filtered to retain only winter-period data in western India. Further details on bird tagging can be found in the methods section of (Baraiya *et al*. 2025). Of these five, one bird was tracked for only one winter season, while the remaining four were tracked for three winter seasons; therefore, data from the four birds that were tracked for three winter seasons were used in this research. Daily movement distance was calculated as the cumulative distance travelled by each individual per day, obtained by summing step lengths between consecutive fixes. To facilitate a direct comparison of movement patterns across winters, all dates were converted into a standardised seasonal index, referred to as Winter Day. Winter Day was defined as a Julian-like variable that began on 1st October (Winter Day = 1) and increased sequentially through the wintering period to 15^th^ March. Dates occurring after 31 December were adjusted to ensure continuity across calendar years. This transformation enabled modelling of intra-seasonal dynamics while maintaining consistency across winters.

GPS locations of the four individuals tracked for three winters were spatially clustered to delineate discrete winter roost sites. Roost sites were defined as discrete wetland water bodies used nocturnally by tracked individuals. A roost site was included in the analysis only if at least one individual used it for at least seven consecutive nights during any of the three study winters. Wetland boundaries were delineated annually in QGIS using Sentinel-2 imagery acquired in December of each study year, representing peak winter inundation extent. Daily movement tracks were classified as cyclic when an individual departed from and returned to the same wetland boundary within a single calendar day, verified spatially in QGIS. Tracks corresponding to inter-site shifts between roost wetlands were identified and excluded. This procedure yielded 1,813 cyclic daily tracks across 13 roost wetlands (Fig. 1, Supplementary Table S1). In order to characterise the habitat composition around each roost site, a 99% Minimum Convex Polygon (MCP) was generated using all associated GPS locations. Each roost-specific MCP was overlaid with a 10 m resolution Sentinel-2 imagery classified by ESRI (Karra *et al*. 2021) for the year 2023 in QGIS to quantify the static proportional composition of major land-use classes, including cropland, rangeland, and built-up areas. In addition to proportional metrics, landscape heterogeneity within each MCP was quantified using the Shannon diversity index and an evenness metric, reflecting habitat diversity and dominance structure, respectively (Supplementary Fig. 1). All spatial metrics were linked to daily movement observations based on the association with roost sites.

**Figure 1.**
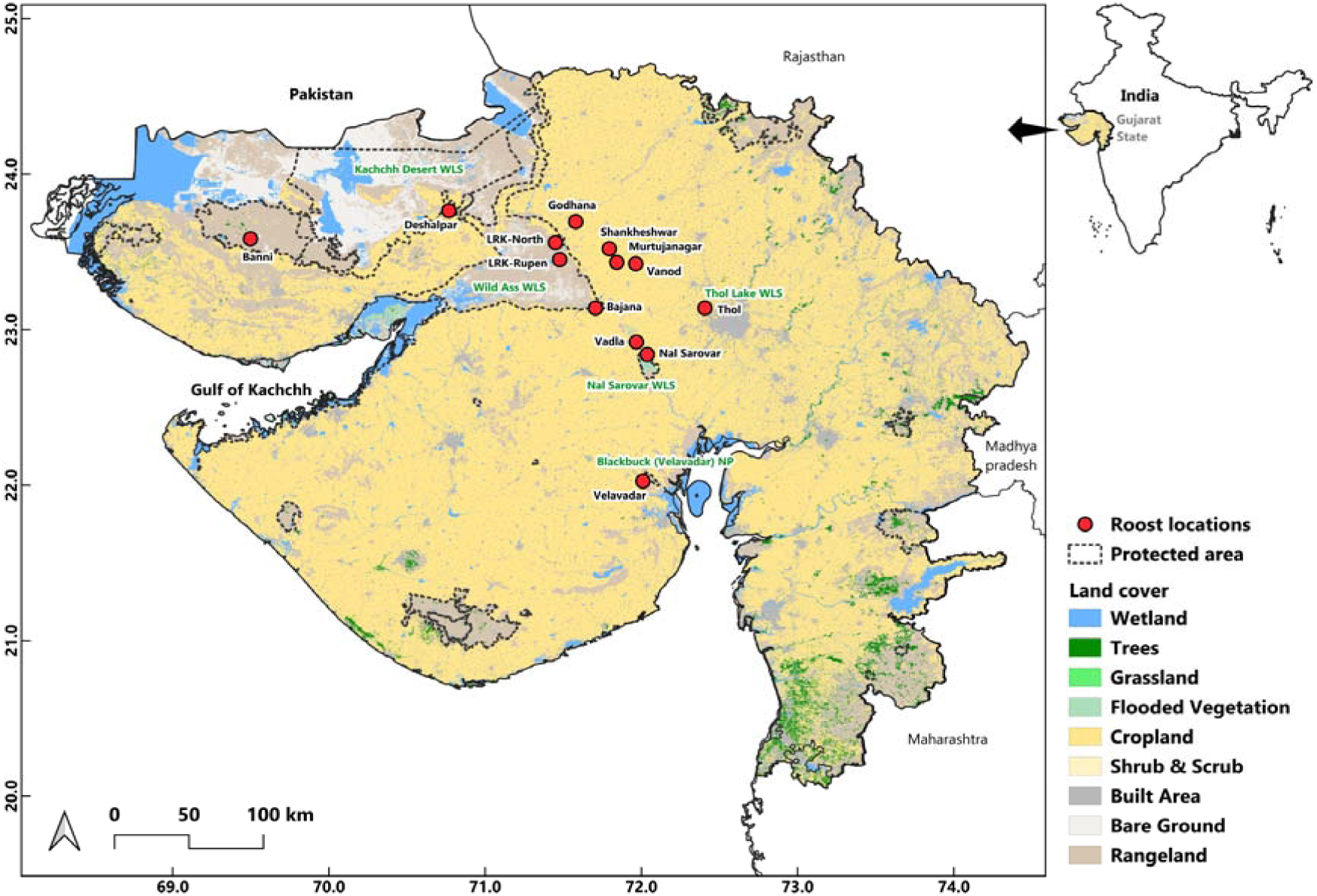
A map showing the identified roost site of the tagged Common Cranes in the State of Gujarat, western India (WLS: Wildlife Sanctuary; NP: National Park)

### Drivers of daily movement

Exploratory analysis indicated that daily movement distances were positively skewed. Consequently, movement distance was modelled using a Gamma distribution with a log link function, appropriate for continuous, non-negative response variables. Collinearity among predictor variables was assessed using pairwise correlation matrices and variance inflation factors derived from linear proxy models using the car package. A strong negative correlation was detected between cropland and rangeland proportions (Supplementary Table S2), and, therefore, predictors were conservatively combined within candidate models to avoid inflated parameter uncertainty. Daily movement distance was modelled using Generalised Additive Mixed Model (GAMM) fitted with the mgcv package (Wood 2017). All models included a smooth term for Winter Day using a thin-plate regression spline to capture nonlinear intra-seasonal trends. Individual identity and winter year were included as random-effect smooths using penalised regression splines to account for repeated measurements and hierarchical structure. Ecological and anthropogenic predictors, including land-use proportions and landscape metrics, were included as parametric fixed effects. A biologically informed candidate model set comprising sixteen models was specified, representing alternative combinations of land-use composition and landscape structure variables. Models were fitted using restricted maximum likelihood, and model selection was based on Akaike’s Information Criterion (AIC). ΔAIC values and Akaike weights were used to rank and identify the most parsimonious model. The best-supported model was evaluated using residual diagnostics, deviance residual distributions, QQ plots, a concurvity assessment, and basis-dimension adequacy checks implemented via diagnostic tools in the mgcv package. Model fit was further assessed using the ratio of deviance to residual degrees of freedom. All the statistical analyses were conducted in R version 4.5.0 (R Core Team 2025).

### Winter utilisation distributions

To quantify winter space use by Common Cranes, we estimated utilisation distribution (UD) with Dynamic Brownian Bridge Movement Models (dBBMMs) in R version 4.5.0. Unlike conventional kernel density estimators, dBBMMs explicitly incorporate the temporal sequence of animal locations and account for movement paths between successive fixes, thereby providing a more realistic representation of space use from GPS telemetry data. GPS tracking data were first screened for erroneous records by removing locations with missing or invalid coordinates. The filtered locations were converted into movement trajectories and projected from geographic coordinates (WGS84) to Universal Transverse Mercator (UTM Zone 43N), allowing movement distances and utilisation distributions to be calculated in metric units. Dynamic Brownian Bridge Movement Models were then fitted using the brownian.bridge.dyn function in the R package move (Kranstauber *et al*. 2024). The model estimates the probability of an individual’s presence between consecutive locations by incorporating both the elapsed time between fixes and the uncertainty in location estimates. A 20 m location error was specified to account for GPS positional uncertainty. Behavioural changes along movement trajectories were identified using a sliding-window approach with a window size of 31 locations and a margin of 11 locations. These parameters allow the variance of Brownian motion to vary dynamically over time, enabling the model to distinguish periods of localised activity from directed movement. Utilisation distributions were calculated on a raster grid with a resolution sufficient to capture fine-scale spatial structure while maintaining computational efficiency.

The resulting utilisation distribution raster represented the relative probability of space use across the wintering area. To facilitate interpretation, the utilisation distribution was converted to cumulative probability volumes and normalised to values between 0 and 1. Isopleths representing the 50%, 75%, and 99% utilization distributions were extracted. The 50% UD was interpreted as the core-use area, the 75% UD as an intermediate-use area, and the 99% UD as the overall extent of space use. The utilisation distribution maps were prepared in QGIS 3.34 (QGIS Development Team 2023).

## RESULTS

### Drivers of daily movement

Daily movement distances of cranes during winter were best explained by the GAMM that included a non-linear smooth of Winter day, random-effect smooths for individual identity and winter year, and linear effects of landscape composition (proportion cropland, proportion built-up area, and habitat evenness; model M13, ΔAIC = 0, Akaike weight = 0.9999) (Supplementary Table S3). Parametric effects indicated that increasing cropland cover was associated with reduced daily distances (β = - 0.330, SE = 0.066, t = -4.99, p < 0.001). In contrast, a greater proportion of built-up area was associated with increased movement distances (β = 3.033, SE = 0.331, t = 9.15, p < 0.001). Habitat evenness also showed a negative relationship with daily distance travelled (β = -0.762, SE = 0.106, t = -7.18, p < 0.001). The smooth term for Winter Day was strongly supported (estimated degrees of freedom (edf) = 9.65, F = 21.60, p < 0.001), reflecting pronounced non-linear seasonal variation in movement. Random-effect smooths indicated substantial variation among individuals (edf = 2.52, F = 6.50, p < 0.001) and between winter years (edf = 1.95, F = 42.96, p < 0.001). The model explained 27.2% of the deviance (adjusted R² = 0.244) (Table 1). Predicted marginal effects and individual-level estimates are presented in Fig. 2.

**Figure 2.**
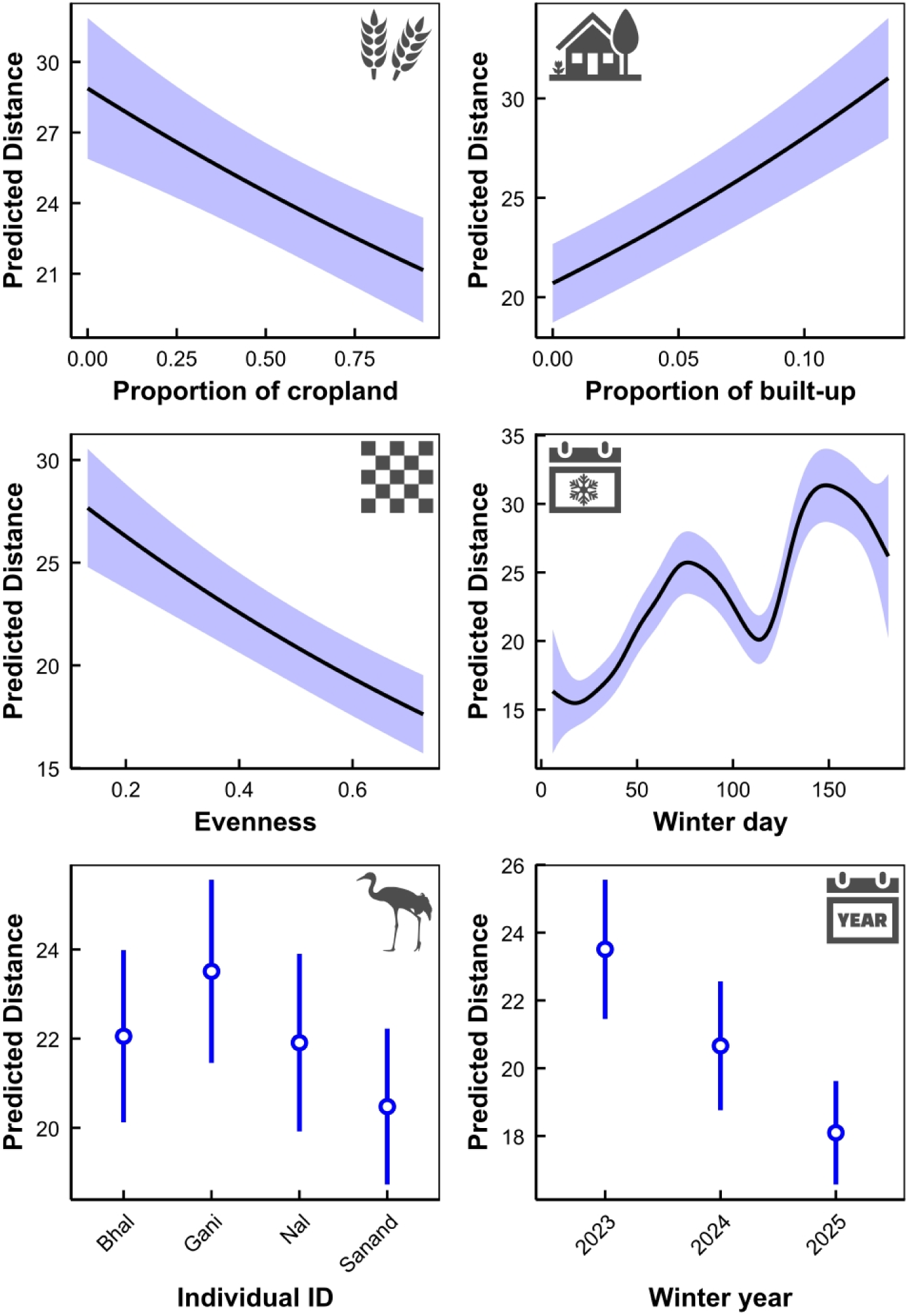
Predicted effects of landscape composition, seasonal progression, and random factors on daily movement distance (km) of the common crane during winter, based on the best-supported GAMM (M13). Lines show predicted distances for continuous predictors with 95% confidence intervals (CI). Dot-whisker plots show the random effects of individual identity and winter year, with dots representing means and whiskers 95% CI, illustrating variation among individuals and across winters.

**Table 1.** Parameter estimates from Generalised Additive Mixed Models (GAMMs) of daily movement distance in the tagged common crane during winter. (A) Baseline GAMM (Best model: M13): population-level effects of cropland, built-up area, and landscape evenness, with seasonal and random effects for individual and winter year.

| Term | Effect type | Estimate ($\beta$) | SE | Test statistic | edf | p-value |
| --- | --- | --- | --- | --- | --- | --- |
| **(A) Baseline GAMM (Best model: M13)** |  |  |  |  |  |  |
| Intercept | Parametric | 3.297 | 0.102 | $t = 32.31$ | – | $< 2.0 \times 10^{-16}$ |
| Cropland proportion | Parametric | –0.330 | 0.066 | $t = -4.99$ | – | $6.75 \times 10^{-6}$ |
| Built-up proportion | Parametric | 3.033 | 0.331 | $t = 9.15$ | – | $< 2.0 \times 10^{-16}$ |
| Landscape evenness | Parametric | –0.762 | 0.106 | $t = -7.18$ | – | $1.03 \times 10^{-12}$ |
| s(WinterDay) | Smooth | – | – | $F = 21.60$ | 9.65 | $< 2.0 \times 10^{-16}$ |
| s(Individual ID) | Random smooth | – | – | $F = 6.50$ | 2.52 | $1.51 \times 10^{-6}$ |
| s(Winter year) | Random smooth | – | – | $F = 42.96$ | 1.95 | $< 2.0 \times 10^{-16}$ |

### Utilisation distributions

Winter space use varied among individuals and years at all utilisation distribution (UD) levels (Figs 2 and 3, Supplementary table S4). Core-use areas (50% UD) ranged from 0.2 to 8 km^2^ across individuals and years. For Bhal, the 50% UD size was 4 km² in 2023, 2 km² in 2024, and 1 km² in 2025, while Gani’s UD area was 8 km^2^, 7 km², and 0.2 km^2^ in 2023, 2024 and 2025, respectively. Nal UD area showed broadly similar 50% UD sizes across winters (4–6 km^2^), whereas Sanand ranged from 1 to 3 km^2^. Intermediate-use areas (75% UD) ranged from 7 to 47 km^2^. Bhal’s 75% UD size gradually decreased over the winters, to 41 km^2^ in 2023, 20 km^2^ in 2024, and 7 km^2^ in 2025. Gani’s UD areas also decreased gradually, to 47 km^2^, 25 km^2^, and 21 km^2^, respectively. Nal remained broadly similar across winters (35, 40 and 35 km^2^), whereas Sanand varied between 31 km², 29 km^2^, and 34 km^2^. The 99% UD showed the widest range of space use, spanning 113–1,068 km^2^. Bhal 99% UD size was 1,068 km^2^ in 2023, 993 km^2^ in 2024 and 113 km^2^ in 2025. Gani recorded 306 km^2^ in 2023, 225 km^2^ in 2024 and 531 km^2^ in 2025. Nal showed relatively consistent 99% UD sizes across winters (566, 591 and 453 km^2^), while Sanand UD ranged from 429 km^2^ in 2023 to 919 km^2^ in 2024 and 786 km^2^ in 2025.

**Figure 2.**
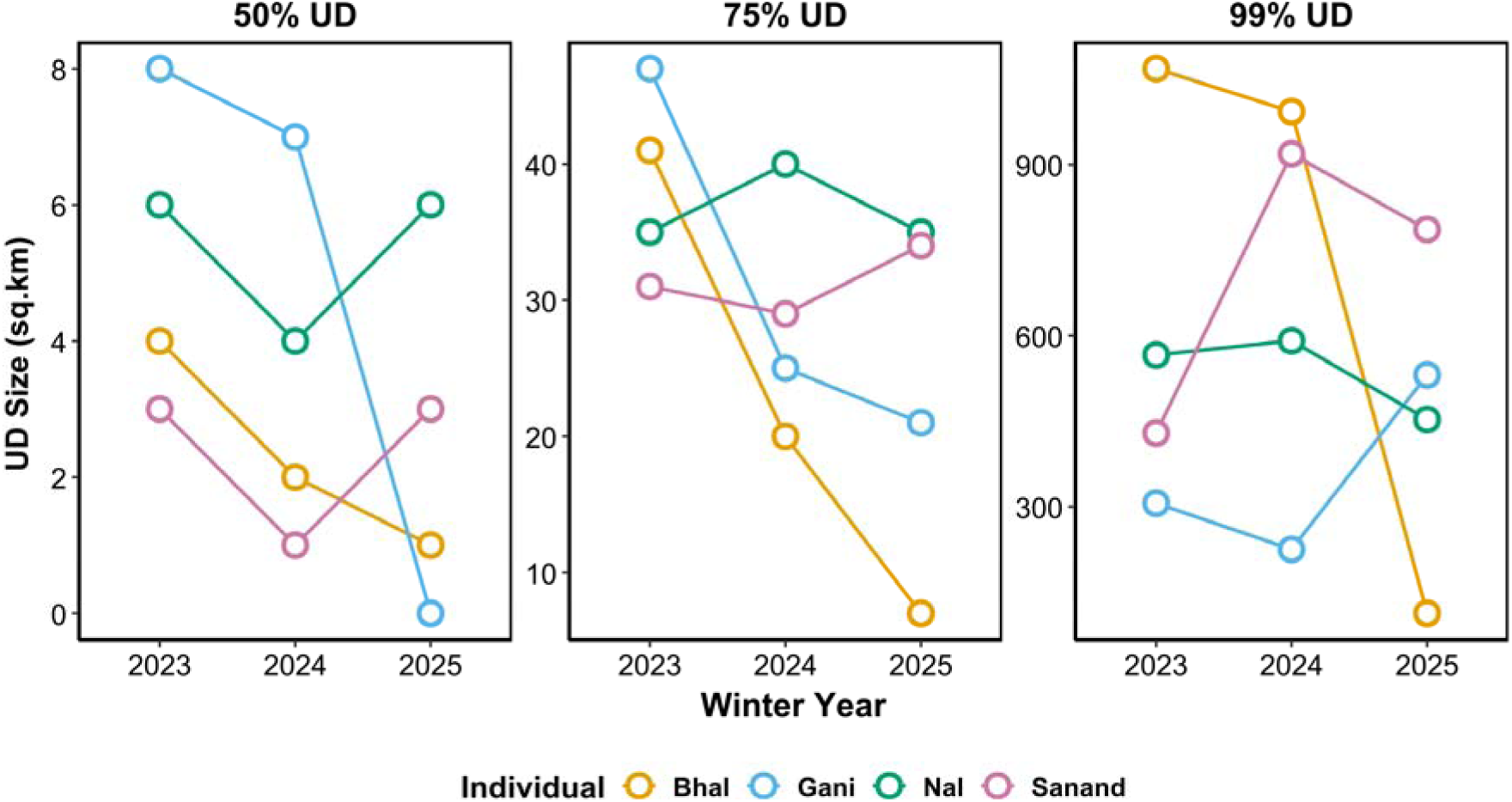
Plots showing inter-individual and inter-annual variation in utilisation distribution (UD) areas across four common cranes over three winter seasons. Colours denote tagged individual names, and circles indicate the estimated area of UD.

**Figure 3.** Utilisation distributions of four GPS-tracked Common Cranes across three wintering seasons (2022-2025) in western India, estimated using the dynamic Brownian Bridge Movement Model (dBBMM). Contours represent 50% (Red), 75% (Green_, and 99% (Blue) utilisation distribution levels, indicating the probability of crane occurrence during the wintering period. (WLS: Wildlife Sanctuary; NP: National Park)

## DISCUSSION

Leveraging GPS tracking data from four individuals over three winters, we present one of the first individual-level assessments of daily movement ecology in wintering Common Cranes across semi-arid agrarian landscapes of western India. We found that in the tagged individuals, (i) daily movement distances exhibited pronounced non-linear seasonal dynamics, (ii) landscape composition, particularly cropland availability, built-up areas, and habitat evenness, shaped daily movement behaviour, and (iii) utilisation distribution extents varied substantially among individuals and across years. Together, these results provide preliminary evidence, based on tracking four individuals over three winters, that wintering cranes may adjust their movement strategies in response to dynamic environmental conditions and heterogeneous human-modified landscapes. Given the small sample size, these interpretations should be treated as exploratory and hypothesis-generating rather than definitive population-level conclusions. Our findings provide preliminary support to the hypothesis that winter movement patterns in the Common Crane are structured by seasonal phenology, landscape composition, and individual behavioural strategies.

### Drivers of daily movement

Our results show that daily movement distances of wintering common cranes may not be random fluctuations but structured responses arising from the interactions among seasonal progression, landscape composition, and individual behavioural tendencies. The observed non-linear temporal trajectory across winter, combined with strong landscape effects and moderate repeatability, indicates that winter displacement reflects adaptive spatial decision-making rather than stochastic variability. The wavy seasonal pattern suggests that daily displacement is governed by shifting patch profitability through winter (Nilsson *et al*. 2018). According to spatial foraging theory, movement distance increases when resource density declines or becomes spatially dispersed, and decreases when resources are abundant and spatially predictable (Van Moorter *et al*. 2013). In agro-wetland systems, harvested croplands initially provide concentrated post-harvest spilt grain, which progressively declines through depletion, decomposition, and farming activity. Early and mid-winter may therefore represent a phase of high patch profitability with reduced search effort, whereas late winter likely requires expanded search to locate remaining high-energy food.

This interpretation is consistent with work showing that cranes adjust environmental niche components rhythmically across seasons, reflecting trade-offs between food, water, and climatic constraints (Yanco *et al*. 2024). Importantly, winter niche use is not static: individuals recalibrate habitat associations as environmental conditions shift. Comparable quadratic seasonal displacement patterns have been reported in whooping cranes, with larger movements early and late in winter (Crouch *et al*. 2024), reinforcing the idea that winter movement follows temporally structured resource dynamics rather than simple directional change. Thus, the seasonal “wave” in our system likely represents a phased adjustment to changing energetic landscapes, in which daily displacement tracks declining agricultural profitability and potentially increasing competition or disturbance.

The strong negative relationship between the proportion of cropland and daily movement distance provides direct behavioural evidence that agricultural landscapes function as high-profitability resource patches during winter. A global synthesis demonstrated that agricultural crops constitute a major component of crane diets worldwide (Hemminger *et al*. 2022). For adaptable species such as the common crane, croplands provide dense, predictable anthropogenic food subsidies that can substantially elevate local carrying capacity. When food is spatially concentrated, optimal foraging predicts reduced movement distances because energy intake per unit search effort increases. Our results are consistent with satellite-based habitat selection, showing strong crane selection for cropland and water bodies in western India (Ram *et al*. 2023) and positive associations with proximity to farmland in China (Duan *et al*. 2025). In such landscapes, daily commuting distances between roost wetlands and feeding sites are minimised, reducing locomotion costs. Crucially, this reduced movement in cropland-dominated landscapes is not merely a matter of habitat preference; it reflects energetic efficiency. Reduced displacement is likely to enhance the net energy balance during winter, a period critical for maintaining body condition before spring migration.

In contrast, built-up areas were associated with increased daily movement. This suggests that urbanisation and infrastructure amplify spatial costs, either through disturbance-mediated avoidance or through fragmentation of foraging patches. Habitat models consistently show crane sensitivity to distances from settlements and roads (Wang *et al*. 2019, Duan *et al*. 2025). Even when cranes exploit agricultural landscapes embedded within human-dominated matrices, urban structures may disrupt direct commuting routes or reduce patch accessibility. Inland wintering Whooping Cranes exhibit larger home ranges and nearly double daily movement distances compared with coastal populations (Crouch *et al*. 2024), illustrating how spatial configuration alters displacement costs. Similarly, anthropogenic environments can reshape winter movement strategies in migratory birds more broadly (Van Doren *et al*. 2021). Thus, increased displacement in built-up landscapes likely reflects compensatory movement: individuals must travel farther to integrate roosting, foraging, and low-disturbance areas. This pattern highlights that anthropogenic landscapes can simultaneously provide food subsidies (cropland) and impose movement penalties (urbanisation), producing a spatial trade-off structure.

The effect of landscape evenness further supports the idea that movement distances reflect spatial integration of resource patches (Borah & Beckman 2023). When landscapes are homogeneous and dominated by cropland, resources are contiguous, allowing efficient exploitation within restricted daily ranges. In heterogeneous mosaics, by contrast, cranes may need to traverse larger distances to connect spatially separated wetlands, fields, and safe resting areas. Studies of common crane staging show gradual spatial shifts that form overlapping “rings” of activity in response to heterogeneous and unpredictable food distribution (Nilsson *et al*. 2018). Such ring-like expansion suggests that movement increases when resource distribution becomes patchier or depleted. Similarly, wintering birds alter their spatial use in response to changes in local habitat profitability (Teitelbaum *et al*. 2023, Schindler *et al*. 2025). Therefore, our landscape configuration results indicate that daily displacement emerges from the spatial arrangement of complementary resources rather than from habitat type alone.

Collectively, our findings suggest that daily displacement during winter reflects dynamic optimisation of energy gain relative to spatial cost. Agricultural landscapes reduce movement by concentrating high-energy resources, whereas urbanisation and heterogeneity increase movement by fragmenting or separating key habitat components. Seasonal waves emerge as profitability shifts across time, and individuals exhibit consistent differences in how they navigate this landscape. This framework aligns with the view that movement is an emergent property of animals continuously balancing energetic gain, predation risk, and disturbance within spatially structured environments (Van Moorter *et al*. 2013). Wintering common cranes in agro-wetland systems, therefore, appear to operate within a subsidy–disturbance trade-off landscape: croplands provide energetic benefits that compress daily displacement, while anthropogenic structures impose spatial costs that expand it. Understanding this dual structure is critical under ongoing agricultural intensification and urban expansion (Hemminger *et al*. 2022). If cropland profitability declines or disturbance intensifies, daily movement distances may increase, potentially elevating energetic expenditure and altering winter body condition. Conversely, concentrated food subsidies may reduce movement but increase dependency on anthropogenic systems.

### Winter utilisation distribution

Winter range distribution size varied substantially both within individuals across winters and among individuals within the same season. This suggests that winter home range size is neither temporally stable within individuals nor spatially uniform across the population. The substantial within-individual variation across winters indicates that seasonal space use is responsive to annual environmental conditions, including variation in the distribution of agricultural resources, hydrology, and disturbance regimes. At the same time, consistent differences among individuals, such as repeatedly expansive ranges in Bhal versus consistently restricted space use in Nal, suggest that cranes differ in the scale at which they integrate winter habitats. Thus, winter home range appears to reflect both environmental contingency and individual-specific spatial strategies.

Compared to previous work in India, our estimates span a much broader range. (Ram *et al*. 2023) reported a mean winter home range of 161.22 ± 172.08 km² based on two tagged individuals monitored during a single season using conventional KDE. In contrast, our estimated utilisation distributions ranged from similarly small areas to several hundred square kilometres. This wider range likely reflects both genuine ecological variability and differences in analytical framework. Our utilisation distributions were derived using dBBMM, which accounts for autocorrelation in telemetry data and provides estimates based on the underlying movement process. In addition, the multi-individual, multi-year design of our study allows assessment of both interannual and inter-individual variability, which could not be evaluated in single-season analyses. Overall, the notable variability observed here indicates that winter range distribution size in common cranes is highly flexible and context-dependent. Seasonal space use can contract to tens of square kilometres under certain conditions or expand to thousands of square kilometres in others, underscoring the importance of multi-year datasets for accurately characterising the spatial ecology of wintering populations.

## CONCLUSION

In conclusion, winter movement ecology of the Common Crane in semi-arid agro-wetland landscapes is predominantly shaped by habitat variability rather than fixed individual differences. Seasonal dynamics followed a nonlinear pattern, consistent with changes in the profitability of agricultural resources throughout winter. Landscape composition further structured movement behaviour: the daily movement distance of the common crane decreased with increasing cropland proportion and increased with increasing built-up area proportion. Winter range distribution size also showed substantial variation both within individuals across years and among individuals within the same season, further reinforcing that seasonal space use is highly context-dependent. Overall, winter spatial behaviour in common cranes emerges as a flexible, environmentally responsive system, underscoring the importance of multi-year, individual-based approaches for understanding movement strategies in dynamic, human-modified landscapes.

## List of abbreviations

AIC: Akaike’s Information Criterion
GAMM: Generalised additive mixed model
LULC: Land-use land-cover
MCP: Maximum convex polygon
NP: National Park
UD: Utilisation distribution
WLS: Wildlife Sanctuary

## Declarations

### Ethics approval and consent to participate

We received permits to capture and tag common cranes from the office of the Principal Chief Conservator of Forest, Gujarat State Forest Department, vide letters WLP/ACF/RTC-3/C/1050-51/2018-19, dated 04-02-2019, and WL/RTC/28/C/852-853/2021-22, dated 19-01-2022.

### Consent for publication

Not applicable

### Availability of data and materials

The datasets used and/or analysed during the current study are available from the corresponding author on reasonable request.

### Authors’ contributions

HLB, RSK, and AB conceived the study. RSK and AB secured funding. HLB and RSK arranged fieldwork logistics. HLB and RSK participated in fieldwork. HLB conducted data analyses. HLB wrote the original draft of the manuscript. All authors reviewed, edited and approved the final manuscript.

## Acknowledgements

The funding support was provided by the PowerGrid Corporation of India Ltd. (PGCIL), and we would like to specifically thank officials of PGCIL - Dr R.K. Srivastava and Mr Suvendu Kumar Kar. We thank the Director and Dean of the Wildlife Institute of India for their support during the study. We thank the Principal Chief Conservator of Forest, Gujarat State Forest Department, for the necessary permissions to carry out bird captures and tagging. We thank Mr Shwetank Pandit (Rtd. IFS) and Mr V.J. Rana (Rtd. IFS), Sh. P. Purushothama (IFS), Dr Chaudhary (IFS), Mr Dipak Solanki (GFS), and Mr Swapnil Patel (GFS) of the Gujarat State Forest Department for their kind support in facilitating fieldwork. We thank Mr Gani Sama and the late Mr Karshan Padhar, the frontline staff of the Nal Sarovar Ramsar Site, for their expert help in trapping birds. We thank Mr Gaurav Sirola, all interns, and field support staff for their support during fieldwork. We thank Mr Shailesh Thakor, caretaker at Thol Ramsar Site, and the people of Vadla village for providing logistical support during fieldwork.

## REFERENCES

Avgar, T., Potts, J.R., Lewis, M.A. & Boyce, M.S. 2016. Integrated step selection analysis: bridging the gap between resource selection and animal movement. Methods in Ecology and Evolution 7: 619–630.

Baraiya, H.L., Sirola, G., Baroth, A. & Kumar, R.S. 2025. Tracking the long way around: seasonal migration strategies, detours and spatial bottlenecks in common cranes wintering in western India. Anim Biotelemetry 13: 38.

Barnosky, A.D., Hadly, E.A., Bascompte, J., Berlow, E.L., Brown, J.H., Fortelius, M., Getz, W.M., Harte, J., Hastings, A., Marquet, P.A., Martinez, N.D., Mooers, A., Roopnarine, P., Vermeij, G., Williams, J.W., Gillespie, R., Kitzes, J., Marshall, C., Matzke, N., Mindell, D.P., Revilla, E. & Smith, A.B. 2012. Approaching a state shift in Earth’s biosphere. Nature 486: 52–58.

Beckmann, M., Gerstner, K., Akin-Fajiye, M., Ceau□u, S., Kambach, S., Kinlock, N.L., Phillips, H.R.P., Verhagen, W., Gurevitch, J., Klotz, S., Newbold, T., Verburg, P.H., Winter, M. & Seppelt, R. 2019. Conventional land-use intensification reduces species richness and increases production: A global meta-analysis. Global Change Biology 25: 1941–1956.

Borah, B. & Beckman, N.G. 2023. Bird movement patterns in an agricultural landscape are mediated by both habitat conditions and traits. Biotropica 55: 1069–1080.

Buitendijk, N.H., de Jager, M., Hornman, M., Kruckenberg, H., Kölzsch, A., Moonen, S. & Nolet, B.A. 2022. More grazing, more damage? Assessed yield loss on agricultural grassland relates nonlinearly to goose grazing pressure. Journal of Applied Ecology 59: 2878–2889.

Butler, S.J., Mattison, E.H.A., Glithero, N.J., Robinson, L.J., Atkinson, P.W., Gillings, S., Vickery, J.A. & Norris, K. 2010. Resource availability and the persistence of seed-eating bird populations in agricultural landscapesL: a mechanistic modelling approach. Journal of Applied Ecology 47: 67–75.

Cao, C. & Song, W. 2022. Progress and prospect of ecological risks of land use change. Front. Environ. Sci. 10.

Clausen, K.K., Dalby, L., Heldbjerg, H., Cao, L. & Fox, A.D. 2024. Using tracking data to assess seasonal habitat use and conflict potential of Greylag Geese in Danish intensive agricultural landscapes. Eur J Wildl Res 71: 6.

Cordes, L.S., Bishop, C.M., Börger, L., Nabe-Nielsen, J. & Harris, S.M. 2025. Optimal movement decisions in complex landscapes. Trends in Ecology & Evolution 40: 960–969.

Crouch, C., Caven, A., Bradshaw, M., Fernald, K., Butler, M. & Kalisek, M. 2024. Space use and movements of inland wintering Whooping Cranes in the Aransas-Wood Buffalo population. ACE 19: art16.

Das, S., Srivastava, A. & Hore, U. 2025. Impact of agricultural land use on diversity and structure of farmland birds. Agriculture, Ecosystems & Environment 381: 109438.

Doherty, T.S., Hays, G.C. & Driscoll, D.A. 2021. Human disturbance causes widespread disruption of animal movement. Nat Ecol Evol 5: 513–519.

Duan, Y., Wang, S., Cao, R., Feng, J., Ge, J. & Wang, T. 2025. Tracking habitat use of migratory birds in a humanLdominated stopover site using deep learning and acoustic indices. Ecosphere 16: e70493.

Fox, A.D., Madsen, J., Boyd, H., Kuijken, E., Norriss, D.W., Tombre, I.M. & Stroud, D.A. 2005. Effects of agricultural change on abundance, fitness components and distribution of two arcticLnesting goose populations. Global Change Biology 11: 881–893.

Haddad, N.M., Brudvig, L.A., Clobert, J., Davies, K.F., Gonzalez, A., Holt, R.D., Lovejoy, T.E., Sexton, J.O., Austin, M.P., Collins, C.D., Cook, W.M., Damschen, E.I., Ewers, R.M., Foster, B.L., Jenkins, C.N., King, A.J., Laurance, W.F., Levey, D.J., Margules, C.R., Melbourne, B.A., Nicholls, A.O., Orrock, J.L., Song, D.-X. & Townshend, J.R. 2015. Habitat fragmentation and its lasting impact on Earth’s ecosystems. Sci. Adv. 1: e1500052.

Hemminger, K., König, H., Månsson, J., Bellingrath□Kimura, S. & Nilsson, L. 2022. Winners and losers of land use change: A systematic review of interactions between the world’s crane species (*Gruidae*) and the agricultural sector. Ecology and Evolution 12: e8719.

Higuchi, H., Javed, S., Nagendran, M. & Fujita, M. 2008. Spring Migration of Eurasian Cranes Grus grus from Gujarat, India to Their Northern Breeding Grounds. Global Environmental Research 12: 69–74.

Jia, Y., Zhang, Y., Lei, J., Jiao, S., Lei, G., Yu, X. & Liu, G. 2019. Activity Patterns of four Cranes in Poyang Lake, China: Indication of Habitat Naturalness. Wetlands 39: 45–53.

Johnsgard, P.A. 1983. Cranes of the World [complete work]. Indiana University Press, Bloomington, Indiana, Bloomington, Indiana.

Karra, K., Kontgis, C., Statman-Weil, Z., Mazzariello, J.C., Mathis, M. & Brumby, S.P. 2021. Global land use/land cover with Sentinel-2 and deep learning. In: 2021 IEEE International Geoscience and Remote Sensing Symposium IGARSS, pp. 4704–4707. IEEE.

Khachar, S., Patankar, H.R., Gaekwad, A., Mundkur, T., Pravez, R. & Naik, R.M. 1987. Wintering Cranes in Gujarat State. In: Proceedings 1987. International Crane Foundation, Baraboo, Wisconsin, USA.

Kong, D., Luo, W., Liu, Q., Li, Z., Huan, G., Zhang, J. & Yang, X. 2018. Habitat use, preference, and utilization distribution of two crane species (Genus: *Grus*) in Huize National Nature Reserve, Yunnan–Guizhou Plateau, China. PeerJ 6: e5105.

Kranstauber, B., Smolla, M. & Scharf, A.K. 2024. move: Visualizing and Analyzing Animal Track Data.

Ming-Ming, Z., Can-Shi, H., Xi-Jiao, S. & Hai-Jun, S. 2021. Seasonal Migration and Daily Movement Patterns of Sympatric Overwintering Black-Necked Cranes (Grus nigricollis) and Common Cranes (Grus grus) in Caohai, Guizhou, China. Waterbirds 44.

Morales, J.M., Moorcroft, P.R., Matthiopoulos, J., Frair, J.L., Kie, J.G., Powell, R.A., Merrill, E.H. & Haydon, D.T. 2010. Building the bridge between animal movement and population dynamics. Philos Trans R Soc Lond B Biol Sci 365: 2289–2301.

Nathan, R., Getz, W.M., Revilla, E., Holyoak, M., Kadmon, R., Saltz, D. & Smouse, P.E. 2008. A movement ecology paradigm for unifying organismal movement research. Proc. Natl. Acad. Sci. U.S.A. 105: 19052–19059.

Nilsson, L., Aronsson, M., Persson, J. & Månsson, J. 2018. Drifting space use of common cranes—Is there a mismatch between daytime behaviour and management? Ecological Indicators 85: 556–562.

Nilsson, L., Bunnefeld, N., Persson, J., Žydelis, R. & Månsson, J. 2019. Conservation success or increased crop damage risk? The Natura 2000 network for a thriving migratory and protected bird. Biological Conservation 236: 1–7.

Nilsson, L., Persson, J., Bunnefeld, N. & Månsson, J. 2020. Central place foraging in a humanLdominated landscape: how do common cranes select feeding sites? Journal of Avian Biology 51: jav.02487.

Overton, C.T. & Casazza, M.L. 2023. Movement behavior, habitat selection, and functional responses to habitat availability among four species of wintering waterfowl in California. Front. Ecol. Evol. 11: 1232704.

Patterson, I.L., Bearhop, S., Shaw, J.M., Hilton, G.M. & McIntosh, A.L.S. 2026. Forage over fear: Assessing drivers of site use in response to shooting disturbance in a managed goose species. Journal of Applied Ecology 63: e70265.

QGIS Development Team. 2023. QGIS Geographic Information System.

R Core Team. 2025. R: A language and environment for statistical computing.

Ram, M., Gadhavi, D., Sahu, A., Srivastava, N., Rather, T.A., Jhala, L., Kapadi, P., Vala, K., Zala, Y., Modi, V., Jhala, D., Patel, A., Baraiya, S. & Devaliya, D. 2023. Satellite Telemetry Insights into the Winter Habitat Use and Movement Ecology of Common and Demoiselle Cranes. Birds 4: 337–358.

Rigal, S., Dakos, V., Alonso, H., Auniņš, A., Benkő, Z., Brotons, L., Chodkiewicz, T., Chylarecki, P., De Carli, E., Del Moral, J.C., Domşa, C., Escandell, V., Fontaine, B., Foppen, R., Gregory, R., Harris, S., Herrando, S., Husby, M., Ieronymidou, C., Jiguet, F., Kennedy, J., Klvaňová, A., Kmecl, P., Kuczyński, L., Kurlavičius, P., Kålås, J.A., Lehikoinen, A., Lindström, Å., Lorrillière, R., Moshøj, C., Nellis, R., Noble, D., Eskildsen, D.P., Paquet, J.-Y., Pélissié, M., Pladevall, C., Portolou, D., Reif, J., Schmid, H., Seaman, B., Szabo, Z.D., Szép, T., Florenzano, G.T., Teufelbauer, N., Trautmann, S., Van Turnhout, C., Vermouzek, Z., Vikstrøm, T., Voříšek, P., Weiserbs, A. & Devictor, V. 2023. Farmland practices are driving bird population decline across Europe. Proc. Natl. Acad. Sci. U.S.A. 120: e2216573120.

Roshier, D.A., Doerr, V.A.J. & Doerr, E.D. 2008. Animal movement in dynamic landscapes: interaction between behavioural strategies and resource distributions. Oecologia 156: 465–477.

Rosin, Z.M., Skórka, P., Wylegała, P., Krąkowski, B., Tobolka, M., Myczko, Ł., Sparks, T.H. & Tryjanowski, P. 2012. Landscape structure, human disturbance and crop management affect foraging ground selection by migrating geese. J Ornithol 153: 747–759.

Roy, P., Ramachandran, R., Ravan, S., Kanawade, V., Sarangi, C. & Thakur, P. 2022. Anthropogenic Land Use and Land Cover Changes-A Review on Its Environmental Consequences and Climate Change. Journal of the Indian Society of Remote Sensing 7: 1–26.

Schindler, A.R., Fox, A.D., Walsh, A.J., Griffin, L.R., Kelly, S.B.A., Cao, L. & Weegman, M.D. 2025. Habitat conditions during winter explain movement among subpopulations of a declining migratory bird. Mov Ecol 13: 74.

Schindler, A.R., Fox, A.D., Wikle, C.K., Ballard, B.M., Walsh, A.J., Kelly, S.B.A., Cao, L., Griffin, L.R. & Weegman, M.D. 2024. Energetic trade-offs in migration decision-making, reproductive effort and subsequent parental care in a long-distance migratory bird. Proc Biol Sci 291.

Soriano-Redondo, A., Franco, A.M.A., Acácio, M., Payo-Payo, A., Martins, B.H., Moreira, F. & Catry, I. 2023. Fitness, behavioral, and energetic trade-offs of different migratory strategies in a partially migratory species. Ecology 104: e4151.

Teitelbaum, C.S., Bachner, N.C. & Hall, R.J. 2023. PostLmigratory nonbreeding movements of birds: A review and case study. Ecology and Evolution 13: e9893.

Teitelbaum, C.S., Prosser, D.J., Ackerman, J.T., Ahmed, S., Alam, A.B.M.S., Azmiri, K.Z., Batbayar, N., Bêty, J., Blake□Bradshaw, A., Boiko, D., Buitendijk, N.H., Buler, J.J., Cabot, D., Casazza, M.L., Cohen, B., Davaasuren, B., Farau, S., Feddersen, J., Fieberg, J., Fiedler, W., Glazov, P., Griffin, L.R., Guillemain, M., Hagy, H., Hardy, M.J., Highway, C., Hoffman, D., Kang, T., Keever, A., Kilburn, J., Kölzsch, A., Kruckenberg, H., Laaksonen, T., Ladman, B.S., Lee, H., Lee, S., Lefebvre, J., Legagneux, P., Linssen, H., Madsen, J., Masto, N.M., McWilliams, S., Mezebish Quinn, T., Mitchell, C., Moreau, A., Müskens, G., Newman, S., Nolet, B.A., Nuijten, R.J.M., Osenkowski, J., Overton, C.T., Piironen, A., Plaquin, B., Ramey, A.M., Rodrigue, J., Rodrigues, D., Schreven, K.H.T., Si, Y., Sullivan, J.D., Takekawa, J., Thomas, P.J., Van Toor, M., Waldenström, J., Williams, C.K., Wolfson, D.W., Xu, F., Brosnan, I.G. & De La Cruz, S.E.W. 2026. Waterfowl Move Less in Heterogeneous and HumanLPopulated Landscapes, With Implications for Spread of Avian Influenza Viruses. Ecology Letters 29: e70265.

Therkildsen, O.R., Balsby, T.J.S., Kjeldsen, J.P., Nielsen, R.D., Bladt, J. & Fox, A.D. 2021. Changes in flight paths of large-bodied birds after construction of large terrestrial wind turbines. Journal of Environmental Management 290: 112647.

Tiwari, J.K. & Rahmani, A.R. 1996. The Common Crane Grus grus and its Habitat in Kutch, Gujarat, India. In: Proceedings of the Salim Ali Centenary Seminar 1996, pp. 26–33. Bombay Natural History Society, Bombay.

Valle, D., Attias, N., Cullen, J.A., Hooten, M.B., Giroux, A., Oliveira-Santos, L.G.R., Desbiez, A.L.J. & Fletcher, R.J. 2024. Bridging the gap between movement data and connectivity analysis using the Time-Explicit Habitat Selection (TEHS) model. Mov Ecol 12: 19.

Van Doren, B.M., Conway, G.J., Phillips, R.J., Evans, G.C., Roberts, G.C.M., Liedvogel, M. & Sheldon, B.C. 2021. Human activity shapes the wintering ecology of a migratory bird. Global Change Biology 27: 2715–2727.

Van Moorter, B., Bunnefeld, N., Panzacchi, M., Rolandsen, C.M., Solberg, E.J. & Sæther, B. 2013. Understanding scales of movement: animals ride waves and ripples of environmental change. Journal of Animal Ecology 82: 770–780.

Vasseur, C., Joannon, A., Aviron, S., Burel, F., Meynard, J.-M. & Baudry, J. 2013. The cropping systems mosaic: How does the hidden heterogeneity of agricultural landscapes drive arthropod populations? Agriculture, Ecosystems & Environment 166: 3–14.

Wang, C., Dong, B., Zhu, M., Huang, H., Cui, Y., Gao, X. & Liu, L. 2019. Habitat selection of wintering cranes (Gruidae) in typical lake wetland in the lower reaches of the Yangtze River, China. Environ Sci Pollut Res 26: 8266–8279.

White, R.J. & Razgour, O. 2020. Emerging zoonotic diseases originating in mammals: a systematic review of effects of anthropogenic land-use change. Mammal Review 50: 336–352.

Wood, S.N. 2017. Generalized Additive Models: An Introduction with R, Second Edition, 2nd edn. Chapman and Hall/CRC, New York.

Wu, D., Hu, C., Zhang, M., Li, Z. & Su, H. 2020. Foraging habitat selection of overwintering Black-necked Cranes in the farming area surrounding the Caohai Wetland, Guizhou Province, China. Avian Res 11: 5.

Yanco, S.W., Oliver, R.Y., Iannarilli, F., Carlson, B.S., Heine, G., Mueller, U., Richter, N., Vorneweg, B., Andryushchenko, Y., Batbayar, N., Dagys, M., Desholm, M., Galtbalt, B., Gavrilov, A.E., Goroshko, O.A., Ilyashenko, E.I., Ilyashenko, V.Y., Månsson, J., Mudrik, E.A., Natsagdorj, T., Nilsson, L., Sherub, S., Skov, H., Sukhbaatar, T., Zydelis, R., Wikelski, M., Jetz, W. & Pokrovsky, I. 2024. Migratory birds modulate niche tradeoffs in rhythm with seasons and life history. Proc. Natl. Acad. Sci. U.S.A. 121: e2316827121.

