## Supplementary for "Space Use and Drivers of Daily Movement in the Common Crane *Grus grus* Wintering in the Agricultural Landscape of Western India"

**Table S1.** Overview of GPS tracking data for individual Common Cranes included in the analyses.

| **Given name of the tagged Common Crane** | **No. of winter seasons tracked** | **No. of GPS fixes** | **No. of tracking days** | **No. of Roost sites used** |
| --- | --- | --- | --- | --- |
| **Bhal** | 3 | 51049 | 450 | 9 |
| **Gani** | 3 | 50898 | 492 | 3 |
| **Nal** | 3 | 48667 | 420 | 5 |
| **Sanand** | 3 | 60379 | 451 | 5 |

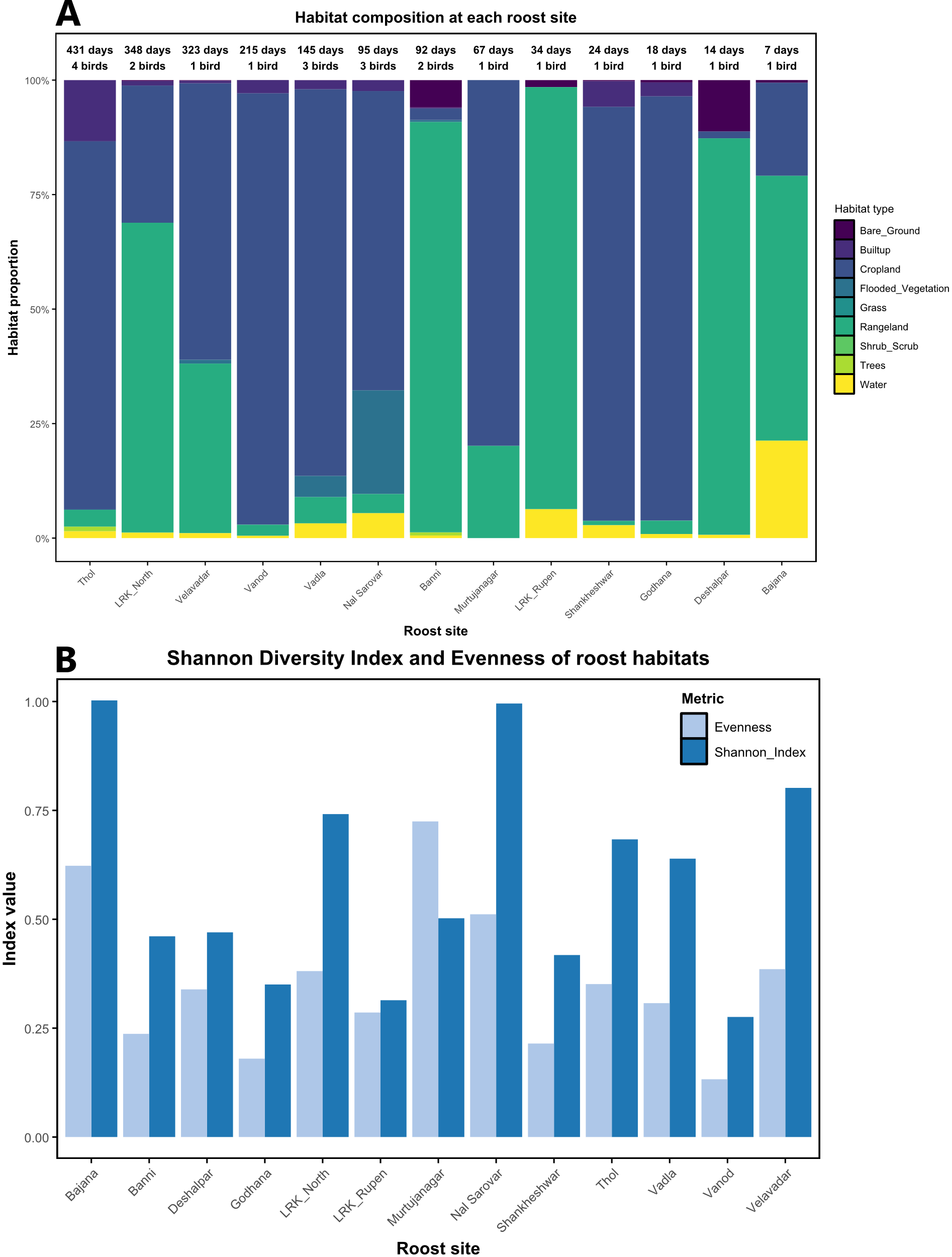

**Figure S1** (A) Stacked bar plot showing the habitat composition surrounding the studied roost sites. The labels above each bar indicate the duration of roost use (days) and the number of tagged individuals that used the respective roost site. (B) Shannon diversity index and habitat evenness of the habitat surrounding the studied roost sites.

**Table S2.** (A) Correlation matrix and (B) variance inflation factors (VIF) for landscape predictors used in GAMM analysis of common crane daily movements.

| (A) Pairwise Pearson correlations among predictors | | | | | |
| --- | --- | --- | --- | --- | --- |
| **Predictor** | **prop_Cropland** | **prop_Rangeland** | **prop_Builtup** | **Shannon_Index** | **Evenness** |
| **prop_Cropland** | 1 | -0.97 | 0.48 | -0.22 | -0.17 |
| **prop_Rangeland** | -0.97 | 1 | -0.58 | 0.11 | 0.11 |
| **prop_Builtup** | 0.48 | -0.58 | 1 | 0.03 | -0.08 |
| **Shannon_Index** | -0.22 | 0.11 | 0.03 | 1 | 0.66 |
| **Evenness** | -0.17 | 0.11 | -0.08 | 0.66 | 1 |

| (B) Variance inflation factors (VIF) from linear proxy model | |
| --- | --- |
| **Predictor** | **VIF** |
| prop_Cropland | 25.76 |
| prop_Rangeland | 28.72 |
| prop_Builtup | 1.91 |
| Shannon_Index | 2.1 |
| Evenness | 1.8 |

**Table S3.** Model selection for candidate GAMMs explaining daily winter movement distances of Common Cranes. All models included a smooth effect of Winter Day (s(WinterDay)) and random effects of individual identity (s(Individual_ID)) and winter year (s(Wint_year)). ΔAIC values are relative to the best-supported model (M13), and Akaike weights indicate relative support among models.

| **Rank** | **Model** | **Habitat predictors included** | **AIC** | **ΔAIC** | **Akaike weight** |
| --- | --- | --- | --- | --- | --- |
| 1 | M13 | Cropland + Built-up + Evenness | 12878.87 | 0 | 9.999985 × 10⁻¹ |
| 2 | M15 | Rangeland + Built-up + Evenness | 12905.68 | 26.81 | 1.508616 × 10⁻⁶ |
| 3 | M12 | Cropland + Built-up + Shannon | 12913.02 | 34.15 | 3.847629 × 10⁻⁸ |
| 4 | M6 | Cropland + Built-up | 12934.9 | 56.03 | 6.808884 × 10⁻¹³ |
| 5 | M14 | Rangeland + Built-up + Shannon | 12942.12 | 63.25 | 1.840603 × 10⁻¹⁴ |
| 6 | M11 | Rangeland + Evenness | 12954.03 | 75.16 | 4.778275 × 10⁻¹⁷ |
| 7 | M7 | Rangeland + Built-up | 12959.87 | 81 | 2.576992 × 10⁻¹⁸ |
| 8 | M3 | Built-up | 12963.62 | 84.75 | 3.959167 × 10⁻¹⁹ |
| 9 | M5 | Evenness | 12965.4 | 86.53 | 1.618744 × 10⁻¹⁹ |
| 10 | M9 | Cropland + Evenness | 12966.24 | 87.37 | 1.067926 × 10⁻¹⁹ |
| 11 | M10 | Rangeland + Shannon | 12993.69 | 114.82 | 1.167988 × 10⁻²⁵ |
| 12 | M4 | Shannon | 13004.98 | 126.12 | 4.114641 × 10⁻²⁸ |
| 13 | M8 | Cropland + Shannon | 13005.33 | 126.46 | 3.457713 × 10⁻²⁸ |
| 14 | M2 | Rangeland | 13032.09 | 153.22 | 5.342883 × 10⁻³⁴ |
| 15 | M0 | – (only WinterDay + random effects) | 13038.51 | 159.64 | 2.161843 × 10⁻³⁵ |
| 16 | M1 | Cropland | 13039.54 | 160.67 | 1.292268 × 10⁻³⁵ |

**Table S4.** A table showing the area of 50%, 75% and 99% utilisation distribution of four tagged Common Cranes across three winter seasons.

| **Given name of the tagged Common Crane** | **50% UD (sq.km)** | | | **75% UD (sq.km)** | | | **99% UD (sq.km)** | | |
| --- | --- | --- | --- | --- | --- | --- | --- | --- | --- |
|  | **2023** | **2024** | **2025** | **2023** | **2024** | **2025** | **2023** | **2024** | **2025** |
| **Bhal** | 4 | 2 | 1 | 41 | 20 | 7 | 1068 | 993 | 113 |
| **Gani** | 8 | 7 | 0.2 | 47 | 25 | 21 | 306 | 225 | 531 |
| **Nal** | 6 | 4 | 6 | 35 | 40 | 35 | 566 | 591 | 453 |
| **Sanand** | 3 | 1 | 3 | 31 | 29 | 34 | 429 | 919 | 786 |
